# The low and high postpubertal ethanol use: long-term effects on drinkers’ reproductive parameters and ethanol-naive offspring development

**DOI:** 10.1101/2021.07.16.452713

**Authors:** Vanessa Caroline Fioravante, Alana Rezende Godoi, Victória Mokarzel de Barros Camargo, Patricia Fernanda Felipe Pinheiro, Marcelo Martinez, Carlos Roberto Padovani, Francisco Eduardo Martinez

## Abstract

The relationship between adolescent ethanol uses and its impacts throughout life is not conclusive. Thus, we evaluated if the low and high consumption of ethanol during the postpuberty period interferes with reproduction in adulthood, the ethanol-naive offspring development and if there are dose-related effects. Females and males’ rats were divided into three groups: low drinker (L), with UChA rats fed with ethanol ad libitum drinking < 1.9 g / kg / day, high drinker (H), with UChB rats fed with ethanol ad libitum drinking from 2 to 5 g / kg / day, and control (C), with rats without access to ethanol. The L and H groups were exposed to ethanol 10% (v/ v) from 65 to 80 days, with withdrawal after this period. The study was conducted in two phases. The retrospective analysis (1st phase) verified the consumption of ethanol between sexes, the litter size, and the sex ratio of offspring. The gestational and reproductive parameters of parents and the development of pups were analyzed in the 2nd phase. We observed a higher consumption of ethanol in females and a reduced litter size in both drinkers’ groups. Body weight gain and gestational feed consumption were lower in L and H. The offspring’s body weight was also lower associated with alteration in landmarks of physical development. The high postpubertal ethanol use accents the impacts on consumers and offspring. The paternal and maternal reproductive organs weight was altered in group H, with an increase in morphologically abnormal sperm. We conclude that low and high post-pubertal alcohol consumption impairs reproductive parameters, even after withdrawal with long-term effects. Ethanol-naive offspring are also harmed, with effects associated with the dose of ethanol.

## Introduction

Ethanol is one of the main abuse drugs ingested worldwide (Rehm et al. 2009, Balddin et al. 2018) and responsible for approximately 5.2% of global deaths (GBD 2018). Alcohol use prevails among the young and adults with more than 76 million people diagnosed with alcohol-related disorders (Ghazali & Patel 2016; Harding et al. 2016). The implications of consumption need to consider the age, the amount ingested and the consumers’ individual characteristics (HHS 2005, Balddin et al. 2018).

Approximately 15% of couples show signs of infertility (Sharlip et al. 2002; Barazani et al., 2014), being the daily habits, including diet and exposure to toxicants, could modulate reproductive health (Asimes et al. 2018). The ethanol is a toxic agent that disturbs the integrity of biochemical and physiological functions and the development of structures involved in reproduction, causing severe damage to the signaling of hypothalamic-pituitary-gonadal / adrenal axes (HPG/HPA) (Wallock-Montelius et al. 2007). This way, alcohol intake can result in female and male reproductive pathologies confirmed in experimental models (Oremosu & Akang 2015, Srivastava et al. 2018) and humans (Eggert et al. 2004, Sansone et al. 2018) since ethanol could lead to lower sperm quality and ovulatory irregularities (Sengupta et al. 2017). Besides, ethanol-induced epigenetic mechanisms can modify the expression pattern of different tissues on drinkers, as well as is the major mechanism related to descendants’ phenotypes alterations (Asimes et al. 2017).

Studies highlight that heavy drinker on gestation can reduce litter size, increase mortality rate, and impair the offspring, however, regarding the sex ratio, the results are inconclusive (Anderson et al. 1978, Cicero et al. 1994, Vaglenova & Petkov 1998, Liang et al. 2014, Gardebjer et al. 2014). Although high alcoholic use has greater effects on reproduction, low to moderate intake has been still under discussion, requiring constant research (Sansone et al. 2018, Barazani et al., 2014). It found in rodents that prepuberty and preconception ethanol exposure can also be harmful to drinkers and their pups, nevertheless, there is no knowledge about the postpuberty period. Thus, studies that aim to verify postpubertal ethanol and its effects on reproduction can help to elucidate the degree of damages of early ethanol intake and the mechanisms behind it. In order to it, we employed the UCh rats, a voluntary ethanol-drinking model derived from original Wistar rats (Mardones & SegoviaRiquelme 1983). This strain represents a special model for understanding the basis of alcoholism-linked characteristics. We hypothesized that the high and low ethanol-drinking on postpuberty negatively influences the parameters of reproduction in adulthood, even after ethanol withdrawal, and affects the ethanol-naive offspring, with dose-related effects. This way, we evaluated whether the low and high ethanol impairs drinkers’ body weight, litter size and sex ratio offspring, the ethanol-naive offspring development, and parents’ reproductive parameters. Part of this study was carried out by a collection of data over ten years, allowing us to assess different generations.

## Material and methods

### Animals and experimental design

The experiments were in accordance with the Ethical Principles in Animal Research and approved by the Bioscience Institute / UNESP Ethical Committee for Animal Research (protocol n° 051/04). Female (171 ± 6.3 g) and male (231 ± 10.7 g) rats (*Rattus norvegicus albinus*) at 55 days old obtained from the Department of Structural and Functional Biology of Botucatu Bioscience Institute / UNESP were used. The rats were divided into three groups: low drinker (L) constituted by UChA rats, high drinker (H) constituted by UChB rats and control rats (C) without access to ethanol. The low and high ethanol-drinking animals were fed with 1:10 (v / v) ethanol *ad libitum* (free choice for water or ethanol) while control animals were fed only water. The ethanol-drinking groups, L and H, were exposed to ethanol for 15 consecutive days for voluntary consumption. Thus, free access of 1:10 (v / v) ethanol solution was provided for them from post-natal day (PND) 65 to 80, corresponding to postpuberty (Picut et al. 2015). Ethanol was withdrawn after this period. The UChA rats consume a low amount of ethanol, drinking from 0.1 to 1.9 g / kg / day, and UChB rats consume a high amount of ethanol, drinking from 2.0 to 6.0 g / kg / day. Only animals with the adequate consumption profile (low and high ethanol consumption) were selected to continue in the experiment (Mardones & Segovia-Riquelmi 1983). After the selection process of drinkers’ animals from L and H groups, they were alcohol withdrawal to mating. Females of C, L, and H groups were mating to males of C, L, and H, respectively, at 100 days old.

The animals were housed in polypropylene cages (32 cm × 40 cm × 18 cm) and maintained under controlled conditions (25 ± 1 °C, humidity 55 ± 5 %, and light from 6 to 18 h) with access to commercial feed and water *ad libitum*. In this study, we employed a voluntary model of ethanol exposure, UCh rats, avoiding the stress associated with forced feeding and providing knowledge about the effects of voluntary ethanol consumption as observed in society (Gapp et al. 2014, Martinez et al. 2016).

This study was conducted in two experimental phases. The 1^st^ one utilized the retrospective analysis to verify the ethanol consumption, consumer body weight, litter size, and sex ratio of offspring from control (C) and low (L) and high (H) drinkers-ethanol groups. The data were collected throughout ten years (2005 -2015), at Anatomy’s Bioterium (IBB/UNESP). Due to significant results from 1st phase, referring to the reproductive capacity of drinkers, we additionally analysis the gestational parameters (body weight and feed and water consumption on gestation), initial development of ethanol-naïve offspring (landmarks of physical development and body weight), and maternal and paternal reproductive parameters (reproductive organs weight and sperm morphology), on 2^nd^ phase.

### 1^st^ experimental phase: retrospective analysis

#### Ethanol consumption of females and males drinkers

The daily ethanol ingestion was determined in males and females (n = 70 / sex / group) from low (L) and high (H) ethanol groups. The average ethanol consumption was obtained from PND 65-80 and measured by (ethanol consumed (ml) / 15 × 100) / body weight (g).

#### Body weight on postpuberty

The average body weight of females and males from C, L and H groups was compared (n = 25 / sex / group). The animals were weighed at PND 80, period referent to end of ethanol consumption after consecutive use.

#### Litter size and sex ratio of offspring

The litter size and sex ratio of offspring were analyzed from C, L and H (n = 110 couples / group). The sex ratio of offspring was verified by the count of females and males at birth since sex dimorphism in neonates is evidenced by the shorter distance between the anus and the genital tubercle of females (Gallavan et al. 1999). Only the first generation (F1) data were considered in this analysis. The long-term reproductive capacity was determined by the litter size from the first and second-generation (F1 and F2, n = 15 couples / generation / group).

### 2^nd^ experimental phase: parameters of parents and offspring

#### Dams’ parameters on gestation

In order to evaluate the evolution of pregnancy, females of C (n = 8), L (n= 8) and H (n = 8) were mated to males of C (n = 8), L (n = 8) and H (n = 8), respectively, at 100 days old at overnight (one female and one male / cage). Vaginal smear was carried out daily in the morning and the first day of pregnancy was considered when spermatozoa were found. After the pregnancy detection, gestational day (GD) 0, the dams were individualized and monitored. Body weight and feed and water consumption on gestation were weekly measured.

#### Offspring body weight and landmarks of physical development

At birth, the offspring were cut to eight pups (four females and four males) per dam. The body weight of offspring was measured on birth from C, L and H (n = 32 sex / group) and the litter body weight (n = 8 / litter / group) was weekly monitored, from PND 1-21, period that includes the neonatal (PND 0 -7), early infantile (PND 8 -14) and late infantile (PND 15 -21) phases. To evaluate the initial physical development, pinna unfolding, hair growth and eye-opening were also daily observed.

#### Parents reproductive organs weight and adiposity index

The females (n = 8 / group) and males (n = 8 / group) from control (C) and high ethanol drinker (H) were weighted and euthanized by CO_2_ inhalation followed by decapitation at PND 150. Females were killed in the estrus phase. The testis, epididymis, ventral prostate, and seminal vesicle (with fluid) in the males, and ovaries and uterus in the females were removed, dissected, and weighed on analytical balance. The relative organs weight was calculated by organ weight (mg) / body weight (g). The adiposity index was also calculated by [(retroperitoneal fat + visceral fat + epididymal/ovarian fat) / final body weight] * 100.

#### Paternal sperm morphology

The volume of ten microliters of semen was obtained from vas deferens of fathers from C and H group, at PND 150. Seminal fluid was placed on the slide, dried at room temperature for 10 minutes and evaluated under phase-contrast microscopy (400X, total magnification). Two hundred sperm per animal were evaluated for head or flagellar defects (Seed, et al. 1996). Anomalies were classified into head anomalies (neither typical nor isolated hook) or tail anomalies (broken or tail headless) and the data were expressed in percentage (Filler 1993).

#### Statistical analysis

The data were analyzed by the software GraphPad Prism ® (version 7, GraphPad Software, San Diego, CA, USA). A one-way ANOVA (parametric data) was used in body weight on postpuberty and physical development of offspring. Post-hoc analysis was performed by Tukey’s multiple comparison test. A Kruskal-Wallis (non-parametric data) was used in litter size and offspring sex ratio. Post-hoc analysis was performed by multiple comparison Dunn’s test. A two-way ANOVA was employed in dams’ parameters on gestation and offspring body weight gain. Post-hoc analysis was performed by Sidak’s multiple comparison test. Time, treatment, and interaction values were expressed in the figure and table legends. Unpaired t test (parametric data) was employed in the ethanol consumption, parents’ reproductive organs weight and adiposity index and paternal sperm morphology. Results were expressed as mean ± S.D. (standard deviation) or median and interquartile range. The differences were considered significant when P < 0.05.

## Results

### 1^st^ experimental phase: retrospective analysis

#### Females consumed more ethanol than males

We observed sex-specific differences related to ethanol consumption in both low (P = 0.0004) and high (P = 0.0383) ethanol groups. Females (L: 1.56 ± 0.25 g / kg / day; H: 4.90 ± 1.89 g / kg / day) ingested higher amount of ethanol than males (L: 1.27 ± 0.34 g / kg / day; H: 4.27± 1.53 g / kg / day).

#### The long-term postpuberty ethanol use decreased body weight

There was a lower body weight in the L and H groups, independently of sex, compared to C (Table 1).

**Table 1.**
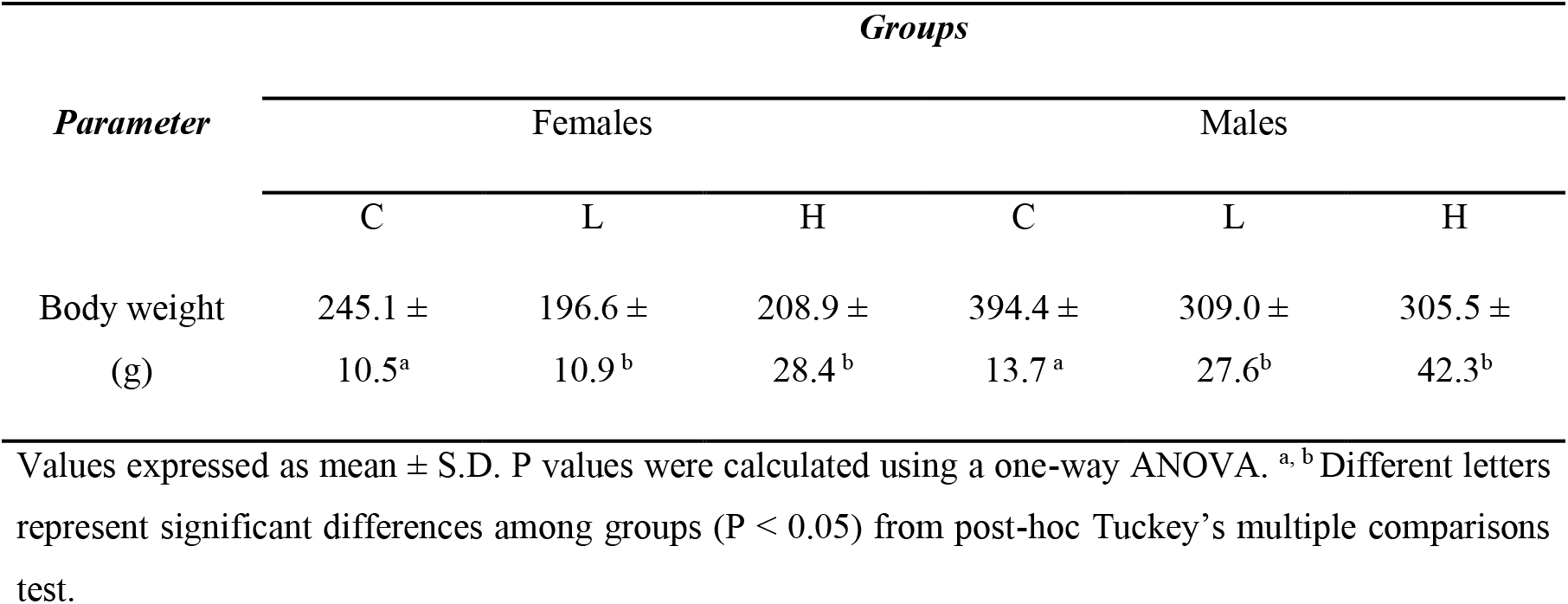
Comparison of females and males body weight from control (C), low (L) and high (H) ethanol groups on post-natal day 80, period correspondent to the end of consecutive ethanol consumption between drinkers’ groups (n = 30 / sex / group).

#### The ethanol uses reduced litter size with dose-related effects, but not impacted on sex ratio of offspring

The litter size from L and H groups was lower compared to C. We observed reduced litter size in the H (Figure 1A) between drinkers’ groups. Thus, the greater ethanol use was the most damaging to the litter size. The comparison of litter size between generations is represented in Figure 1B. Only the H group showed reduced litter size comparing F1 to the F2.

**Figure 1.**
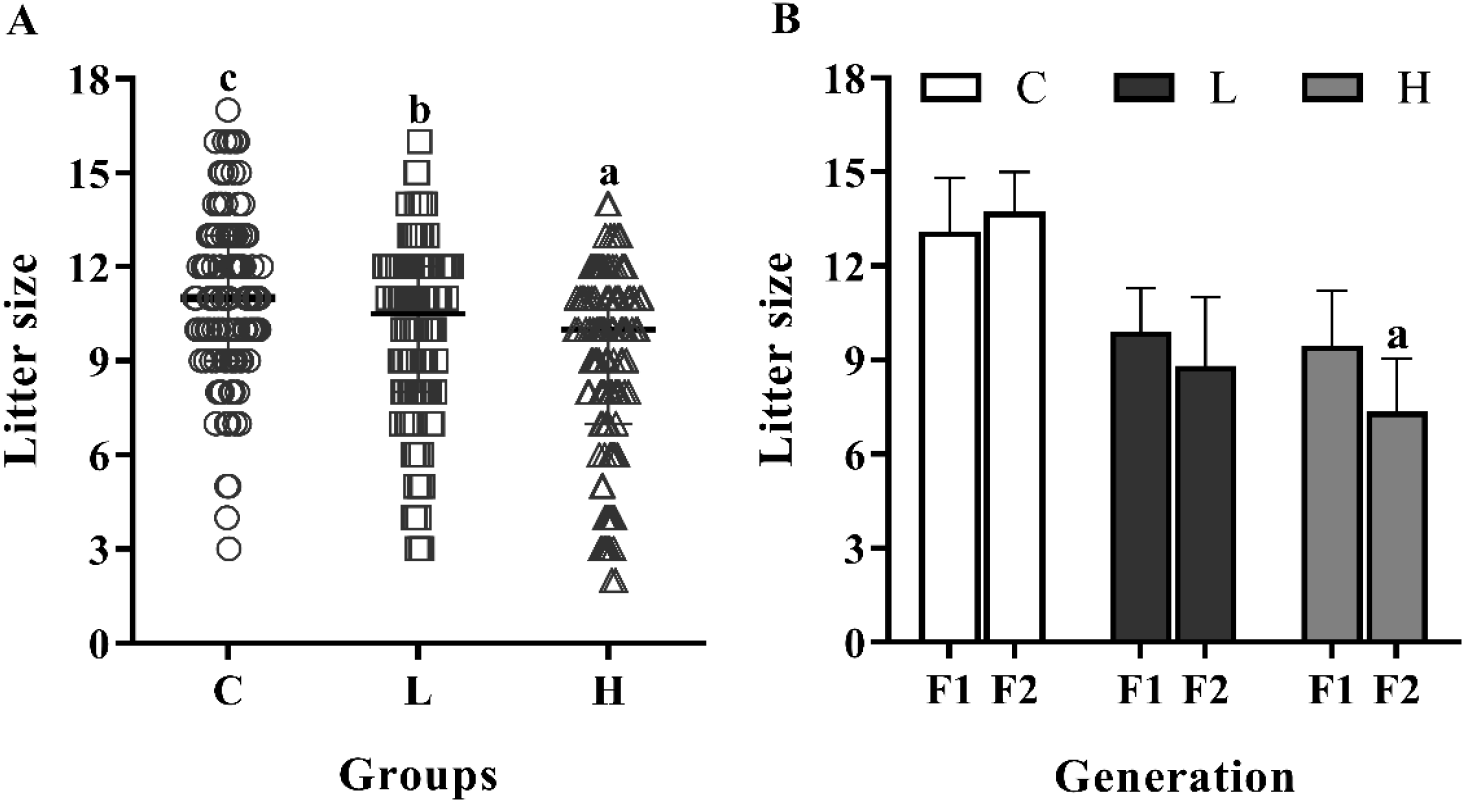
Comparison of litter size from control (C), low drinker (L), and high drinker (H) groups. A) Litter size from C (n = 110), L (n = 110) and H (n = 110) groups. Values expressed as median and interquartile range. P values were calculated using a Kruskal-Wallis test. ^a, b, c^ Different letters represent significant differences among groups (P < 0.05) from post hoc Dunn’s multiple comparisons test. B) Litter size of first (F1) and second (F2) generation from C (n = 15), L (n = 15) and H (n = 15) groups. Values expressed as mean ± S.D. P values were calculated using a two-way ANOVA. ^a^ Significant difference between generations (P < 0.05) from post-hoc Sidak’s multiple comparison test. Figure 1B: P_Inter_ = 0.0595, P_Time_ = 0.0709, P_Treat_ < 0.0001.

There were no differences in the sex ratio of offspring from C, L and H groups (females: C = 51.38 % ± 15.82; L = 50.53 % ± 18.02; H = 50.74 % ± 17.30; males: C = 48.62 % ± 15.53; L = 49.47 % ± 18.02; H = 49.26 % ± 17.00).

### 2^nd^ experimental phase: parameters of parents and offspring

#### The low and high ethanol use decreased gestational body weight gain and feed consumption, no dose-related effects

The maternal body weight on GD 0 and GD 21 was lower in both postpubertal ethanol exposed groups compared to control (GD0 C: 285.4 ± 14.9, L: 236.7 ± 11.5, H: 246.4 ± 11.8; GD 21 C: 315.0 ± 34.7, L: 286.7 ± 12.7, H: 287.2 ± 7.8). However, the dams body weight gain from L and H groups was lower only on the 3^rd^ gestational week (Figure 2C). Regarding the feed and water consumption, we observed lower consumption in the L and H dams on the 2^nd^ and 3^rd^ gestational weeks, respectively (Figures 2 A -B).

**Figure 2.**
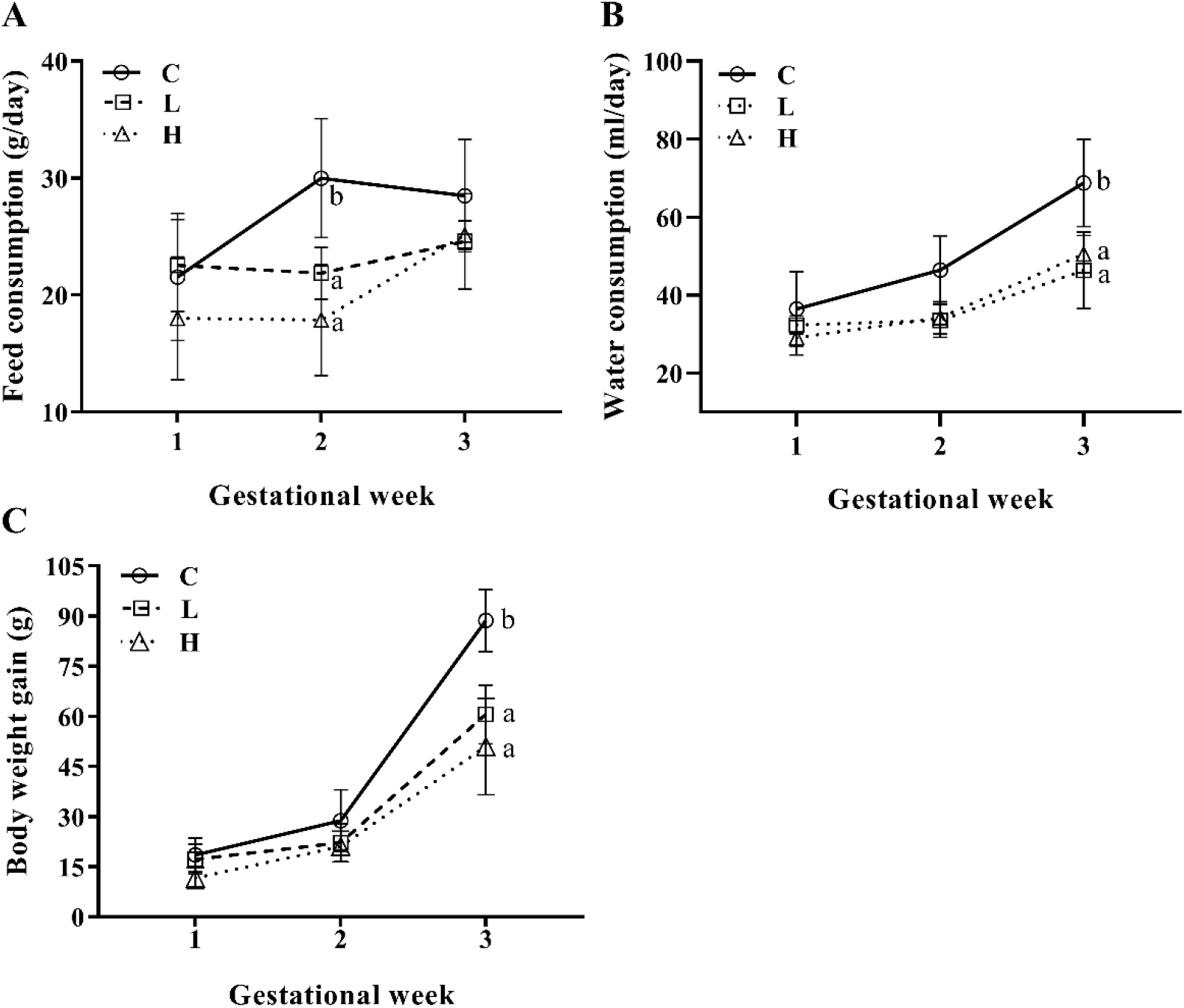
Dams’ parameters on gestation of control (C), low drinker (L) and high drinker (H). A) Feed consumption, B) Water consumption, C) Body weight gain of C (n = 8), L (n = 8) and H (n = 8). Values expressed as mean ± S.D. P values were calculated using a two-way ANOVA. ^a, b^ Different letters represent significant differences among groups (p < 0.05) from post-hoc Sidak’s multiple comparison test. Figures 2A: P_Inter_ = 0.0122, P_Time_ < 0.0008, P_Treat_ < 0.0001; 2B: P _Inter_ = 0.0701, P_Time_ < 0.0001, P_Treat_ < 0.0001; 2C: P_Inter_ < 0.0001, P_Time_ < 0.0001, P_Treat_ < 0.0001.

#### Low and high postpubertal parental ethanol use impaired body weight and physical development of ethanol-naive offspring, with dose-related effects

Figure 3 represents the parameters of female and male offspring from C, L, and H groups. The pups sired by low and high postpubertal parental ethanol use had a lower body weight at birth and throughout the infant period (PND 1 -21) compared to control (Figure 3 A). The damages on the offspring were correlated to the amount of ethanol consumed by parents since the offspring from high drinkers’ group showed lower body weight compared to offspring from low drinkers’ group.

**Figure 3.**
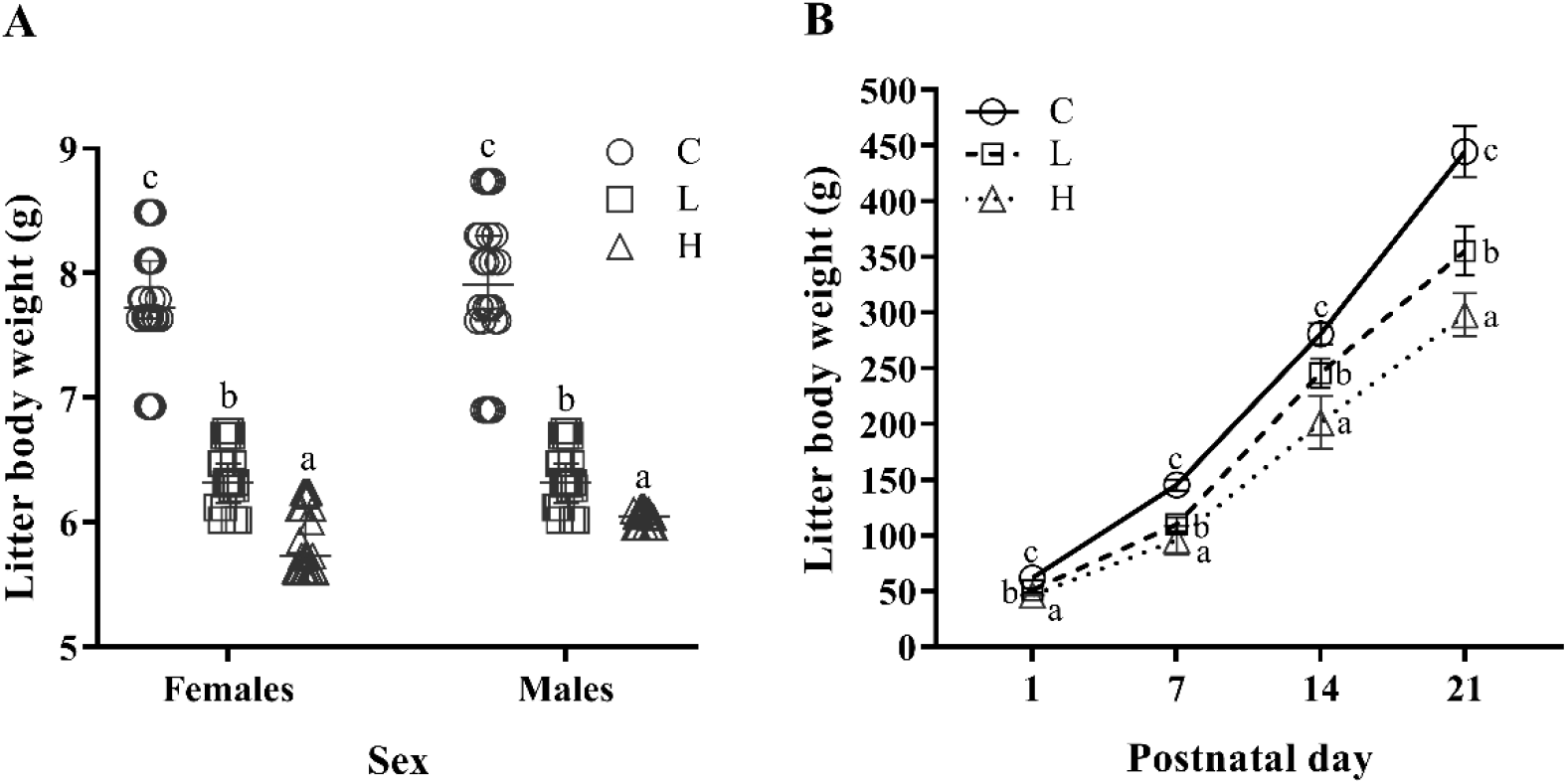
Parameters of female and male offspring from control (C), low drinker (L) and high drinker (H) groups. A) Body weight on birth of females and males (n = 32 / sex / group). Values expressed as median and interquartile range. P values were calculated using a Kruskal-Wallis test. ^a, b, c^ Different letters represent significant differences among groups (P < 0.05) from post hoc Dunn’s multiple comparisons test. B) Litter body weight (females and males) on postnatal day 1 to 21 (n = 8 /litter/ group). Values expressed as mean ± S.D. P values were calculated using a two-way ANOVA. ^a, b, c^ Different letters represent significant differences among groups (P < 0.05) from post-hoc Sidak’s multiple comparison test. Figure 3B: P_Inter_ < 0.0001, P_Time_ < 0.0001, P_Treat_ < 0.0001.

The landmarks of physical development were changed on the offspring from low and high drinkers’ groups. It was observed earlier eye-opening in the offspring from L and delayed hair growth in the H (Table 2).

**Table 2.** Comparison of the mean day of physical development landmarks in days in the offspring (n = 8 / litter / group) from control (C), low drinker (L) and high drinker (H) groups.

| *Parameters*<br>(days) | *Groups* |  |  |
| --- | --- | --- | --- |
|  | **C** | **L** | **H** |
| Pinna unfolding | 2.5 ± 0.5 | 2.3 ± 0.8 | 3.1 ± 0.8 |
| Hair growth | 4.7 ± 0.4 <sup>a</sup> | 4.8 ± 0.5 <sup>a</sup> | 5.7 ± 0.4 <sup>b</sup> |
| Eye-opening | 14.2 ± 0.3 <sup>b</sup> | 13.6 ± 0.3 <sup>a</sup> | 14.3 ± 0.5 <sup>b</sup> |

Values expressed as mean ± S.D. P values were calculated using a one-way ANOVA. <sup>a, b</sup> Different letters represent significant differences among groups (P < 0.05) from post-hoc Tukey's multiple comparison test.

#### High postpubertal ethanol use decreased female and male reproductive organs weight on adulthood and impaired paternal sperm morphology

We did not get data from the parental reproductive organs and the paternal sperm morphology of the low drinker group. Therefore, in this analysis only data from control and high drinkers’ groups were compared.

There was lower uterine weight and adiposity index in the females from H group, but the body weight did not alter. On the other hand, the body weight was lower and sex glands (epididymis and seminal vesicle) was greater in the males from H (Table 3).

**Table 3.** Comparison of body weight, relative reproductive organs weight and adiposity index at post-natal day 150 in the females and males from control (C) and high drinkers (H) groups (n = 8 / sex / group).

| *Parameters* | *Groups* |  |
| --- | --- | --- |
|  | **C** | **H** |
| *Females* |  |  |
| Body weight (g) | 315.01 ± 30.99 | 287.2 ± 8.77 |
| Ovaries (mg/g) | 0.46 ± 0.04 | 0.43 ± 0.02 |
| Uterus (mg/g) | 2.13 ± 0.12 | 1.62 ± 0.15 * |
| Adiposity index | 3.99 ± 0.73 | 3.00 ± 0.31 * |

| *Males* |  |  |
| --- | --- | --- |
| Body weight (g) | 506.05 ± 12.22 | 408.00 ± 16.37 * |
| Testis (mg/g) | 3.67 ± 0.29 | 4.02 ± 0.29 |
| Epididymis (mg/g) | 1.50 ± 0.09 | 1.72 ± 0.02 * |
| Ventral prostate (mg/g) | 3.08 ± 0.22 | 3.28 ± 0.67 |
| Seminal vesicle (mg/g) | 3.95 ± 0.95 | 5.62 ± 0.77 * |
| Adiposity index | 2.93 ± 0.62 | 2.85 ± 0.46 |

Values expressed as mean ± S.D. P values were calculated using a t test. \* Significant difference between groups (P < 0.05).

Regarding paternal sperm morphology, there was an increase in the percentage of sperm with morphologic abnormalities in the early high exposed to ethanol group, including a higher incidence of head and tail defects (Table 5).

**Table 5.** Paternal sperm morphology on post-natal day 150 in the males from control (C) and high drinkers (H) groups (n = 8 / group).

| *Parameters (%)* | *Groups* |  |
| --- | --- | --- |
|  | **C** | **H** |
| Normal sperm | 91.25 (90.38 – 92.63) | 75 (70.38 – 81.25) * |
| Abnormal sperm | 8.75 (7.37 – 9.62) | 25 (18.75 – 29.63) * |
| Broken tail | 0.56 (0.50 – 0.71) | 0.75 (0.00 – 3.00) |
| Headless | 1.56 (0.87 – 2.09) | 4.00 (2.87 – 4.12) * |
| Isolated head | 5.87 (4.50 – 6.87) | 19.75 (12.50 – 23.75) * |

Values expressed as median and interquartile range (IQ1 - IQ3). P values were calculated using a t test. \* Significant difference between groups (P < 0.05).

## Discussion

This is the first study to conduct a retrospective analysis of postpubertal ethanol use and its effects on the reproduction in the voluntary model of low and high ethanol consumption. Our results highlight how lifestyle after adolescence modulates long-term reproductive parameters and future generations. The low and high postpuberty ethanol use impairs reproduction, even after withdrawal, and the development of ethanol-naive offspring, being high alcohol drinking the most harmful.

Ethanol consumption is historically higher in men, however, the difference between sexes has been disappearing (Becker & Koob 2016). We observed greater ethanol ingestion in females from both drinking groups. These findings can be partially attributed to biological differences established by hormonal status during development since estrogen receptors are found in structures belonging to mesocorticolimbic reward pathways and in serotoninergic function (Bethea et al. 2002, Witt 2007, Becker & Koob 2016). Ethanol is a caloric substance and could interfere in regular nutrition (Gruchow et al. 1985), thus, we observed lower body weight in females and males L and H after 15 consecutive days of exposure. Increased energy expenditure and lipid oxidation and poor nutritional status due to a decrease in non-alcoholic calorie intake may be related (Addolorato et al. 1998, Ojeda et al. 2009, Sebastiani et al. 2018).

In this perspective, postpubertal alcohol use also led to lower weight gain and feed consumption on gestation in the L and H dams, with dose-related effects. Increased plasma leptin during abstinence (Kiefer et al. 2005) contributes to decreased food intake. On the other hand, the inefficiency in the absorption of ingested calories can also act, since the metabolism of ethanol changes the intestinal permeability and harms the organic systems function (Bishehsari et al. 2017). In addition, the lower weight gain could be associated with the weight of the pregnant uterus (Kind et al. 2006), fetal and placental (Brett et al. 2014), as we found lower litter size and body weight at birth of the offspring from L and H groups.

Considering the direct-ethanol toxicity, gestational exposure can reduce litter size, especially in the high drinkers (Cicero et al. 1994, Ojeda et al. 2009, Li et al. 2012). Interestingly, we found that postpubertal ethanol use also decreased litter size but did not alter offspring sex ratio. Impacts on blastocyst implantation, oxidative damage to germline DNA compromising the embryo cells, abnormal fetal development, and increased rates of resorption and abortion may be mechanisms contributing to these results (Cicero et al. 1994, Ward et al. 1996, Emanuele et al. 2001, Jana et al. 2010, Jensen et al. 2014). We hypothesize the reproductive capacity of high drinkers was chronically affected, as there was a decreased litter size between the first-and-second-generation.

The reproductive organs’ weight has been used to evaluate the toxicity of the reproductive system (Clegg et al. 2001). We found alterations in the female and male reproductive organs in the high drinkers’ group. Studies noticed an irregularity in the estrous cycle, reduction of LH and FSH, follicular atresia, and damage in uterine endometrial cells in drinkers, with effects related to the dose consumed (Chuffa et al. 2009, Martinez et al. 2016). Acetaldehyde and oxidative stress increase and impairs to HPG/HPA axes are mechanisms that change reproductive hormones balance and, consequently, uterine and ovarian tissues (Buthet et al. 2003, Rachdaoui & Sarkar 2017). We suggest that lower uterine weight in H females could be related to a hormonal imbalance with damage to uterine structure and function. The lower adiposity index observed in females from the H group could also highlight possible malnutrition due to loss of muscle or fat mass (Dasarathy 2016). Relating to males, studies have evaluated alcohol exerting a direct effect on both testosterone metabolism and spermatogenesis (Sansone et al. 2018). In contrast to the literature that reports atrophy of reproductive organs in drinkers (Martinez et al. 2000; 2001), we found an increase in the epididymis and seminal vesicle weight. Alcohol consumption can affect the autonomic nervous system, with sacral denervation leading to erectile dysfunction (Johnson et al. 1986, Julian et al. 2019). Changes in contractility and innervation of these organs increase epididymis sperm reserve and drive to accumulation or retention of fluid in the seminal vesicle (Fernandez et al. 2008). Furthermore, we found an increased percentage of sperm with morphological abnormalities similar to clinical (Pajarinen et al. 1996, La Vignera et al. 2012, Sansone et al. 2018) and experimental (Jana et al. 2010) studies. These increased abnormalities can be either due to failures in the spermatogenic process or in sperm maturation. Inadequate signalization of epididymal factors which plays a role in maturation or low testosterone levels can drive these abnormalities (Koch et al. 2015, Zi et al. 2015). The abnormal testosterone/estradiol ratio has been also associated with decreased semen parameters as well as harm on accessory sex glands (Ramasamy et al. 2016). The lower body weight of H males validates the compromised growth. This parameter can indicate estrogen imbalance, as body weight has been used to measure estrogen potency (Heywood & Wadsworth 1980, Hart 1990). The damage to reproductive organs and testosterone can be observed even after alcoholic withdrawal (Candido et al. 2007), then, we hypothesized toxicity to the HPG/HPA axes, leading to inadequate functioning of accessory sex glands, hormonal imbalance, and reduced sperm quality. Although additional analysis is needed to validate the real harm of ethanol on reproduction, our data indicate that high post-pubertal alcohol use can highly impair long-term reproductive function. Low doses are also harmful, nevertheless, their impacts are lesser than the high doses (Patra et al. 2011; Rahimipour et al. 2012).

Studies highlight preconception ethanol exposure can influence the descendants’ phenotype. In this way, we observed that in addition to the impairment on the gestational parameters of dams early exposed to the low and high ethanol, the offspring also showed reduced body weight, with dose-related effects. Reducing the gestational sac weight, placental efficiency, and function due to maternal and paternal ethanol use (Chang et al. 2017, Gardebjer et al. 2014) could explain partially our results. Preconception nutrition, body weight index, gestational weight gain and food consumption also influence maternal metabolic response in pregnant and fetal outcomes (Kind et al. 2006, Brett et al. 2014). Associated, the paternal experiences play an additional role (Robertson 2005, Rando & Simmons 2015) since the seminal fluid stimulates the female reproductive tract to produce growth factors and cytokines which protect the embryo (Robertson 2005) and changes in the seminal signalization are capable of influencing descendants (Bromfield et al. 2014). The landmarks of physical development on offspring sired by alcoholic use by parents were altered in our study, similar to results by Fioravante et al. (2021). The insulin-like growth factor (IGF) is important for fetal and postnatal development and IGF deficiency implicates signaling pathways and normal body growth (Kanaka-Gantenbein et al. 2003). Similarly, the endothelial growth factor (EGF) plays a role in regulating the activity of epidermal and epithelial tissues as eye-opening and hair growth (Smart et al. 1989, Calamandrei & Alleva 1989). The alterations in the postnatal development of offspring can be also associated with maternal care (Amorim et al., 2011) since alcohol withdrawal accentuates depressive behaviors and reduced time spent on nursing (Pang et al. 201, Workman et al. 2015). We hypothesized a possible endocrine and metabolic programming of the offspring, with the parental ethanol dose as decisive in the course of this programming.

In summary, the results presented here suggest the potentially alarming possibility in which exposure to alcohol on post puberty produces long-term effects, even in withdrawal of alcohol. The ethanol use decreased body weight, gestational feed intake, and litter size. In addition, postpuberty alcohol use increased the percentage of morphologically abnormal sperm and alternated female and male reproductive organs weight. Besides the impairments on consumers, the offspring development and growth were also affected, however, we cannot distinguish which parent contributed the most to observed changes. However, our previous laboratory studies along with published data about preconception maternal ethanol exposure strongly suggest a maternal influence as the main. The parameters evaluated show a dose-effect relationship.

Despite the limitations of this study, we highly believe that the post-adolescent period also acts as a susceptibility window. Future studies are needed to identify the long-term effects of consumption on drinkers’ reproduction even in withdrawal as well as to understand the mechanisms responsible for ethanol-naive offspring outcomes. Possibly, the effects are associated with epigenetic germline modifications, metabolism activity and HPG/HPA axis.

## Funding

This study was financed by the Grant 2018/12354-5, São Paulo Research Foundation (FAPESP), and Coordination for the Improvement of Higher Education Personnel -Brazil (CAPES) -Finance Code 001.

## Notes

### Competing Interest Statement

The authors have declared no competing interest.

